# Brain Age Prediction: Cortical and Subcortical Shape Covariation in the Developing Human Brain

**DOI:** 10.1101/570333

**Authors:** Yihong Zhao, Arno Klein, F. Xavier Castellanos, Michael P. Milham

## Abstract

Cortical development is characterized by distinct spatial and temporal patterns of maturational changes across various cortical shape measures. There is a growing interest in summarizing complex developmental patterns into a single index, which can be used to characterize an individual’s brain age. We conducted this study with two primary aims. First, we sought to quantify covariation patterns for a variety of cortical shape measures, including cortical thickness, gray matter volume, surface area, mean curvature, and travel depth, as well as white matter volume, and subcortical gray matter volume. We examined these measures in a sample of 869 participants aged 5-18 from the Healthy Brain Network (HBN) neurodevelopmental cohort using the Joint and Individual Variation Explained (Lock et al., 2013) method. We validated our results in an independent dataset from the Nathan Kline Institute - Rockland Sample (NKI-RS; N=210) and found remarkable consistency for some covariation patterns. Second, we assessed whether covariation patterns in the brain can be used to accurately predict a person’s chronological age. Using ridge regression, we showed that covariation patterns can predict chronological age with high accuracy, reflected by our ability to cross-validate our model in an independent sample with a correlation coefficient of 0.84 between chronologic and predicted age. These covariation patterns also predicted sex with high accuracy (AUC=0.85), and explained a substantial portion of variation in full scale intelligence quotient (R^2^=0.10). In summary, we found significant covariation across different cortical shape measures and subcortical gray matter volumes. In addition, each shape measure exhibited distinct covariations that could not be accounted for by other shape measures. These covariation patterns accurately predicted chronological age, sex and general cognitive ability. In a subset of NKI-RS, test-retest (<1 month apart, N=120) and longitudinal scans (1.22 ± 0.29 years apart, N=77) were available, allowing us to demonstrate high reliability for the prediction models obtained and the ability to detect subtle differences in the longitudinal scan interval among participants (median and median absolute deviation of absolute differences between predicted age difference and real age difference = 0.53 ± 0.47 years, r=0.24, p-value=0.04).

## 1. Introduction

The human brain undergoes dramatic structural changes throughout the lifespan, including brain maturation processes from childhood to adolescence and young adulthood. Characterizing typical developmental trajectories of brain structure maturation is essential for understanding neurodevelopmental disorders and vulnerabilities of the brain (Gogtay et al., 2004; Toga et al., 2006). Most studies of brain development have focused on the cortex and subcortical structures. For instance, developmental trajectories of cortical thickness, cortical and subcortical gray matter (GM) volume, surface area, mean curvature, and white matter (WM) development in different age groups have been extensively examined using both cross-sectional (Asato et al., 2010; Barnea-Goraly et al., 2005; Hill et al., 2010) and longitudinal data (Giedd et al., 1999; Gogtay et al., 2004; Lenroot et al., 2007; Mensen et al., 2017; Sowell et al., 2004). It is well recognized that cortical development is spatially heterogeneous with individual brain regions following distinct temporal patterns of maturational changes (Toga et al., 2006). To best characterize the complexity of cortical development across different brain regions, investigators have begun to use brain structural features to derive a comprehensive index of brain development (Cole and Franke, 2017; Erus et al., 2015).

Brain-based age prediction is thought to be important for understanding normal brain developmental process in healthy individuals, as well as atypical brain structural/functional developmental patterns that might have clinical implications (Davatzikos et al., 2009; Dosenbach et al., 2010). Among brain structural features, cortical thickness (Khundrakpam et al., 2015; Lewis et al., 2018) and GM volume (Franke et al., 2010) have been used to predict brain age. An increasingly broad range of possible measures exist for characterizing brain cortical structures, including surface area, mean curvature, travel depth, and WM volume; additionally, some studies have used the GM/WM difference or contrast (Franke et al., 2012; Lewis et al., 2018) or developed composite metrics incorporating multiple features to increase the brain age prediction accuracy (Brown et al., 2012; Erus et al., 2015; Lewis et al., 2018). However, it remains unclear to what extent different cortical shape measures covary and whether their covariation patterns are related to chronological age.

To characterize covariation patterns across different cortical and subcortical measures, we use the Joint and Individual Variation Explained (JIVE) method (Lock et al., 2013) to estimate shared (i.e., covariation patterns common across all shape measures) and distinct (i.e., covariation patterns specific to an individual shape measure) covariation patterns across six cortical shape measures (cortical thickness, GM volume, surface area, mean curvature, travel depth, WM volume) and subcortical GM volumes. JIVE was developed for simultaneously identifying consistent patterns across multiple data types and patterns unique to individual data types. Specifically, JIVE decomposes the total variance in data into three terms: joint variation across all seven shape measures, structured variation unique to individual shape measures, and residual noise that should be discarded from analyses. JIVE has been used in cancer studies to identify genetic variants across different data platforms (e.g., genotyping, mRNA and miRNA expression) (Hellton and Thoresen, 2016; O’Connell and Lock, 2016) and to identify the common variance between task-based fMRI connectivity and a wide range of behavioral measures (Yu et al., 2017). We hypothesized that integrative analysis of different cortical and subcortical shape measures can provide a more comprehensive picture of brain cortical and subcortical development from childhood to early adulthood, and thus may generate insights into the relationship between brain maturation and cognitive development.

The main purposes of this study were to investigate (1) whether there are common and distinct covariation patterns across a set of the aforementioned six brain cortical shape measures along with subcortical GM volume, and (2) whether those common and distinct covariation patterns can accurately predict age using ridge regression (Hoerl and Kennard, 1970). Due to the impact of sex on brain structural differences and the relationships between cognitive function and brain structure (Erus et al., 2015; Kaczkurkin et al., 2019; Ruigrok et al., 2014; Schmithorst, 2009), we also assessed whether those JIVE components could predict sex and full scale intelligence quotient (FSIQ). To determine whether this approach is generalizable, we validated both our JIVE analysis and age prediction results in an independent dataset. Importantly, this replication involved the application of the JIVE components and prediction models derived in one sample (i.e., CMI Healthy Brain Network) to an independent sample that employed distinct imaging protocols and recruitment strategies.

## 2. Materials and methods

### 2.1 Datasets

Detailed descriptions about the participants and neuroimaging data used in the study are outlined below. All brain features used in this study were extracted from T1-weighted (MPRAGE) scans.

#### 2.1.1 Training data: The Healthy Brain Network cohort

To characterize covariations common across and/or distinct within different structural shape measures, we used T1-weighted MRI scans from the Healthy Brain Network (HBN) project (Alexander et al., 2017). The HBN is an ongoing initiative focused on creating and sharing a biobank comprised of data from up to 10,000 New York City area children and adolescents (https://healthybrainnetwork.org/). A self-referred community sampling strategy is used, with more than 80% of children being identified with one or more mental health or learning disorders. Our analyses were based on N = 869 individuals (530 male, 339 female, mean age = 10.54 ± 3.26, age range: 5.02 – 17.95 years; mean FSIQ: 98.81 ± 16.42; FSIQ range: 50 - 147) collected from three imaging sites used by the HBN. Structural MRIs were acquired at either 1.5T (mobile MRI Scanner in Staten Island, N=630) or 3T (Siemens Tim Trio at Rutgers University, N=11; Siemens Prisma at Cornell Weill Medical College, N=228) using standard T1-weighted sequences. Participants with serious neurological disorders (e.g., Huntington’s disease, amyotrophic lateral sclerosis, multiple sclerosis, cerebral palsy), acute brain dysfunction, diagnosis of schizophrenia, schizoaffective disorder, bipolar disorder, manic or psychotic episode within the past six months, or history of lifetime substance dependence requiring chemical replacement therapy were excluded from the study.

#### 2.1.2 Independent validation data: the NKI-RS longitudinal sample

T1-weighted MRI scans from the Nathan Kline Institute - Rockland Sample (NKI-RS) (Nooner et al., 2012) child longitudinal study (http://rocklandsample.org/child-longitudinal-study) were used as our independent validation data. The child longitudinal study is an ongoing project aimed to generate and share a large-scale, community-ascertained longitudinal sample for understanding comprehensive growth curves of brain function and structure. Per study protocol, T1-weighted MPRAGE scans (TR = 1900; voxel size = 1mm isotropic) are collected for each participant at up to three time points. We had access to the baseline data of N = 210 participants (122 male, 88 female, mean age = 12.31 ± 3.06, age range: 6.68 – 17.94 years; mean FSIQ: 105.00 ± 13.49; FSIQ range: 77 - 142). Recruitment of study participants was designed to maximize community representativeness. Among those with the baseline data, 120 subjects had retest imaging data obtained 3~4 weeks later, and 77 subjects had longitudinal follow-up scans (mean inter-scan interval 1.22 ± 0.29, range 0.31 – 1.92 years).

### 2.2 Brain feature extraction and morphometry

#### 2.2.1 MRI preprocessing

MRI preprocessing entailed two complementary processes: standard MRI preprocessing by the FreeSurfer (Dale et al., 1999) software package (https://surfer.nmr.mgh.harvard.edu) with subsequent feature extraction and shape analysis by the Mindboggle software package (Klein et al., 2017) (https://mindboggle.info). Briefly, FreeSurfer’s recon-all pipeline (v.5.3.0) performed motion correction, intensity normalization, skull stripping, subcortical segmentation, tissue classification, and surface extraction. Preprocessing quality was assessed by a number of standard quality control measurements (Shehzad et al., 2015). MRI data failing quality control analyses were removed from further analysis. The preprocessed T1-weighted MRI data were then imported into Mindboggle to extract shape measures for further analysis. The technical details of Mindboggle have been described elsewhere (Klein et al., 2017). Mindboggle has been shown to improve brain region labeling and provides a variety of precise brain shape estimates.

#### 2.2.2 Brain shape measures

We aimed to characterize covariations among different shape measures for cortical surface and subcortical regions. We focused on six measures, including cortical mean curvature, white matter volume, cortical thickness, surface area, gray matter volume, and travel depth for 62 regions of interest (ROIs) (31 ROIs per hemisphere) per the Desikan–Killiany–Tourville (DKT) protocol (Klein and Tourville, 2012). For a given ROI, we used the median shape value across all vertices in that ROI. Additionally, we included gray matter volume estimates for 16 subcortical ROIs. Thus, the shape measures in this study can be regarded as data from seven different sources.

### 2.3 Data analysis

#### 2.3.1 Dimension reduction by Joint and Independent Variance Explained (JIVE)

Investigating each brain shape individually might fail to identify important interrelationships among different shape measures. We hypothesized the existence of hidden covariation patterns shared across and/or specific to individual shape measures. We therefore used JIVE (Lock et al., 2013) to identify such hidden patterns that might be of biological interest. JIVE extends Principal Component Analysis to data from multiple sources (e.g., different shape measures) and was developed primarily for discovering shared covariation patterns (i.e., joint components) in different sets of measures, as well as systematic variations (i.e., individual components) specific to an individual data source. Specifically, we denote each shape measure by *X*_1_, *X*_2_,…, *X*_7_, where *X_i_, i* = 1,..,7 is a matrix with each row representing a shape measure for a single ROI and each column representing a participant. Since different shape measures have different biological implications and they are at different scales, we regard those as data from different sources. Following Lock et al. (Lock et al., 2013), JIVE decomposes total variation in data into three major parts (joint components, individual components, and error terms):

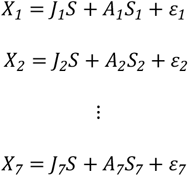

where *S*:*r* × *n* and 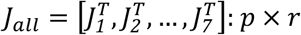 are the joint component scores and loading matrices that capture variations common across seven different shape measures; *S_i_*: *r_i_* × *n* and *A_i_*: *p_i_* × *r_i_* are the individual component scores and loadings suggesting variation specific to individual shape measures; and *ε_i_*: *p_i_* × *n* are error terms. Here *n* is the sample size, 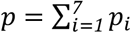, and *p_i_* indicates the total number of ROIs assigned to each shape measure. For a given number of joint components *r* and individual components *r_i_*, the component loadings are obtained by minimizing the overall sum of squared errors 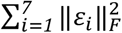, where 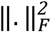 defines the Frobenius norm. The optimal number of joint components *r* and the number of individual components *r_i_s* in this study were determined by a permutation test (Lock et al., 2013) with the significance level set to 0.0001 with 1,000 permutations. All procedures were implemented using the R package ‘r.jive’.

#### 2.3.2 Age, sex and IQ prediction by ridge regression

We used ridge regression models (Hoerl and Kennard, 1970) to assess whether the joint and/or individual components are predictive of age, sex, or FSIQ. Specifically, using JIVE joint and/or individual components as input data, ridge regression models were used to predict chronological age and FSIQ, while ridge logistic regression models were used to predict sex. As a control, we also considered the predictive power of total volumes of subcortical gray matter, intracranium, brain stem, and CSF, which are known to be related to age and sex. Therefore, the input predictors took seven different formats: total volumes only, joint components only, individual components only, total volumes + joint components, total volumes + individual components, joint components + individual components, and total volumes + joint components + individual components.

Briefly, ridge regression models are linear models with ridge penalty which has been shown to offer good prediction performance with high dimensional correlated input data. In mathematic format, given a continuous response variable *y_i_*, and a set of predictors *Z_ij_*, ridge regression estimates the parameters *β_j_s* by minimizing 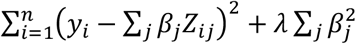. The tuning parameter *λ* controls the model’s complexity. If *λ* = 0, ridge regression becomes a traditional linear regression model. The optimal choice of the *λ* parameter in this study was based on 10-fold cross validation.

#### 2.3.3 Assessment of prediction accuracy

Age and FSIQ prediction accuracy were assessed using two criteria: Mean Absolute Error (MAE) and the coefficient of determination (R^2^), where R^2^ is the proportion of the variance in the dependent variable explained by the model. To determine whether adding additional predictors would improve model prediction accuracy, we report the adjusted R^2^ (i.e., R^2^ adjusted for the number of predictors in the model) for the training model based on HBN data. Sex prediction accuracy was assessed using area under receiver operating characteristic curve (AUC). To demonstrate how the models generalize to data from another study, we predicted age, FSIQ, and sex in the NKI-RS with models trained on HBN data. We report model prediction accuracy assessments from both HBN and NKI-RS data.

## 3. Results

### 3.1 Variance decompositions of shape measures across brain regions

JIVE analysis of 388 brain features (6 cortical shape measures for each of 62 ROIs plus 16 subcortical GM volumes) from the HBN data resulted in a total of 35 lower dimensional representations of brain features (i.e., components). These included 2 joint components, 7, 4, 6, 5, 6, 2, and 3 individual components specific to mean curvature, WM volume, cortical thickness, surface area, GM volume, travel depth, and subcortical GM volumes, respectively. Figure 1 shows the amount of shared and individual variations among the seven shape measures. Overall, joint components were responsible for more variation in cortical WM volume (48.3%), GM volume (52.1%), surface area (45.2%), and subcortical GM volume (41.6%) than in cortical thickness (31.7%), travel depth (15.1%), and mean curvature (11.2%). Covariation between cortical GM volume and surface area are expected as GM volume roughly equals the product of surface area and cortical thickness. In addition, each shape measure has a certain amount of structured variation (ranging from 9.8% to 33.0%) that is unrelated to other shape measures. It should also be noted that we found considerable residual noise in travel depth (67.2%), mean curvature (56.4%), cortical thickness (47.6%), and surface area (42.3%).

**Figure 1:**
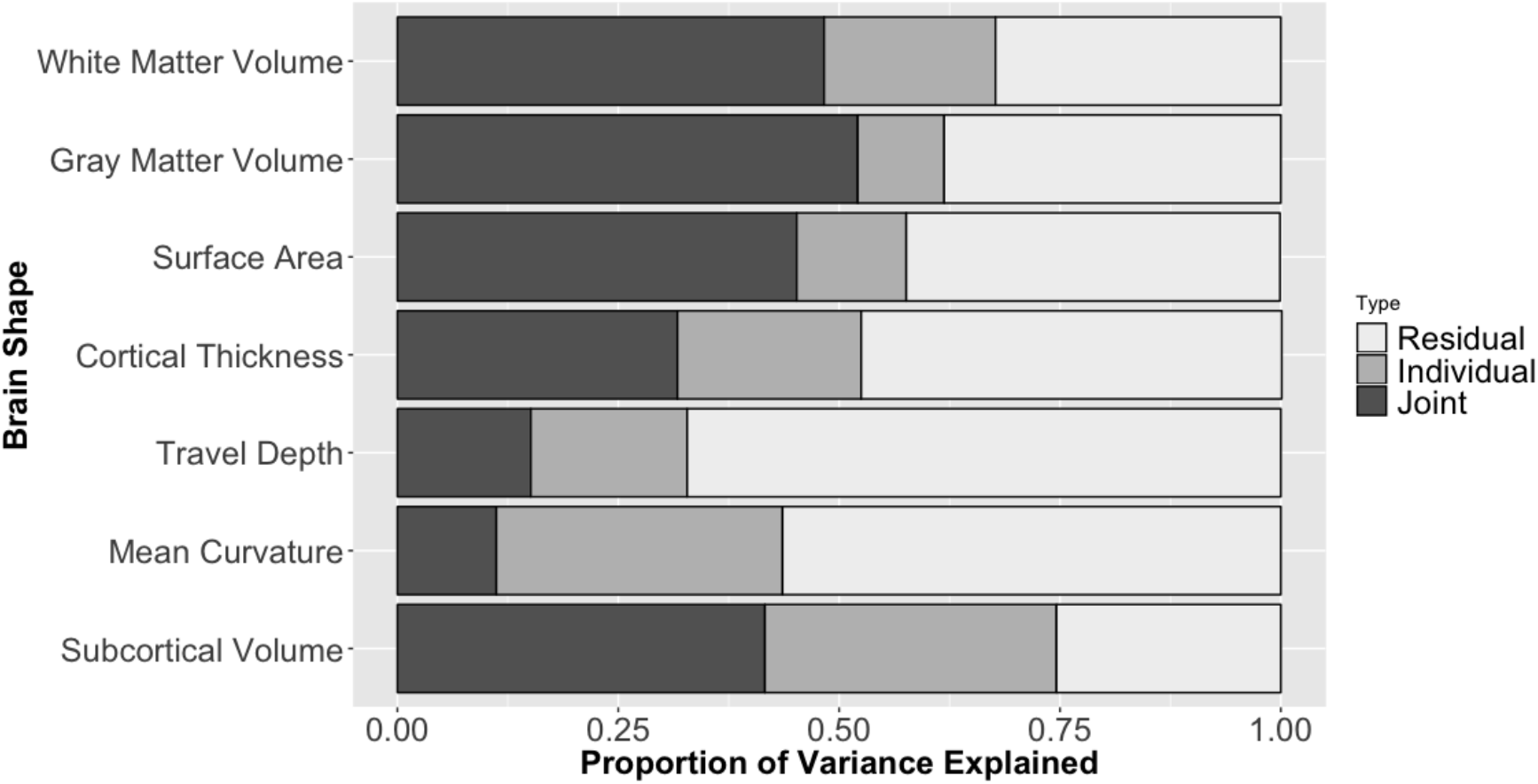
Proportion of variance explained by JIVE components

Of particular interest, we showed that the identified JIVE components were reproducible in an independent dataset. Figure 2 shows the pairwise correlation coefficients between the component loadings from HBN data and those from NKI-RS. The correlation coefficient represents the maximal correlation between an HBN component and its corresponding NKI-RS component. Among the 35 JIVE components, 20 (57.1%) components had correlation coefficients of at least 0.5 between HBN and NKI-RS loadings, and 27 (77.1%) had coefficients of 0.3 or higher. Remarkably, the two joint components along with two travel depth components (individual components specific to travel depth) had correlation coefficients of at least 0.95. Additionally, single individual components specific to WM volume (r=0.92), subcortical component (r=0.87), mean curvature (r=0.86), cortical thickness (r=0.85), surface area (r=0.78) and gray matter volume (r=0.78), respectively, had correlation coefficients above 0.8 or just below (0.78). Thus, a total of 10 JIVE components replicated well in the NKI-RS dataset. It should be noted that most of the individual components specific to cortical GM volume, cortical thickness, and surface area were less consistent between samples, as evidenced by their relatively weak correlations (r < 0.5) in loadings between HBN and NKI-RS components.

**Figure 2:**
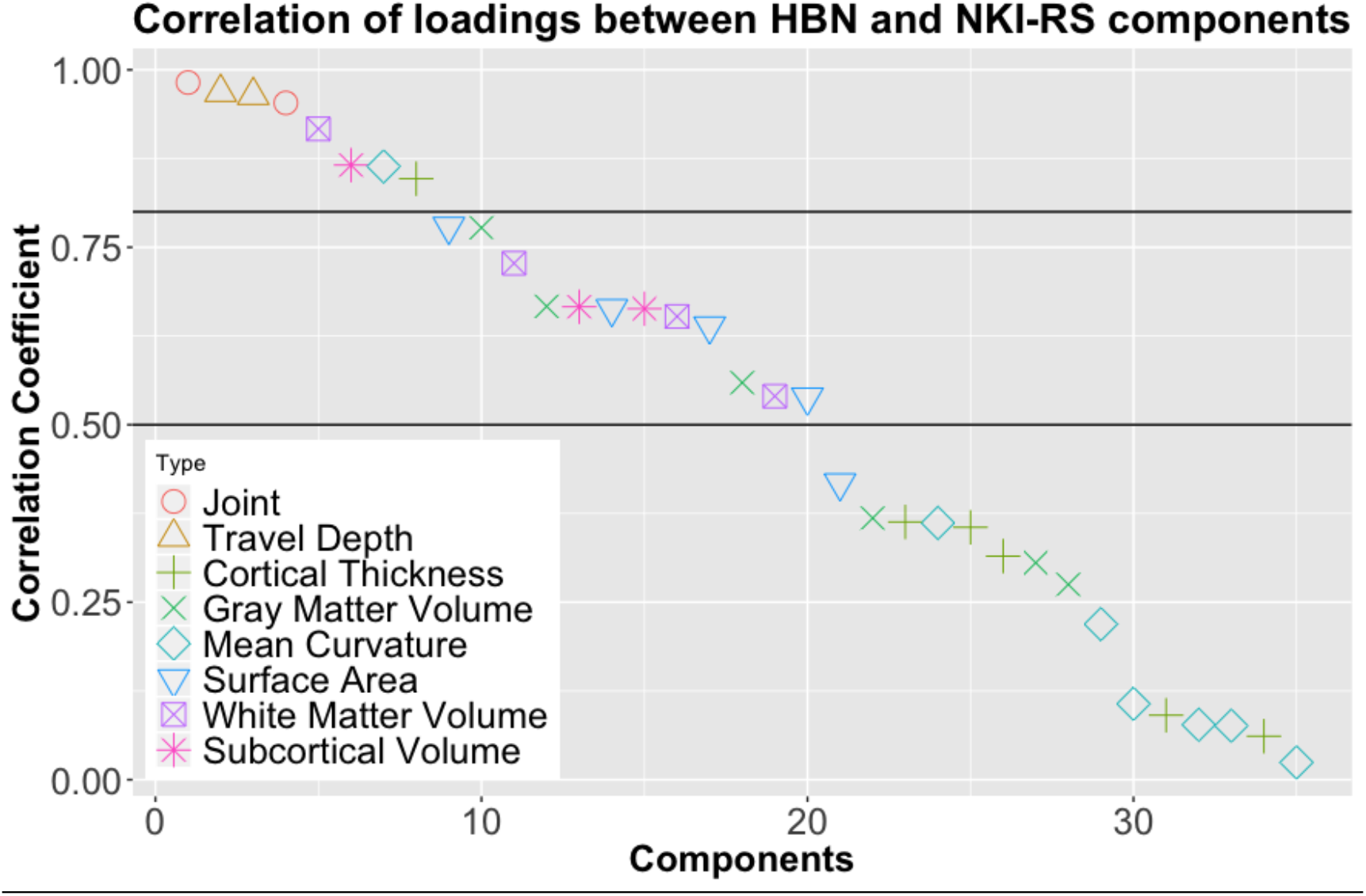
Pairwise correlation coefficients of JIVE component loadings between HBN and NKI-RS

To assess whether the 10 components that replicated in the NKI-RS dataset were related to age, sex, and FSIQ, we employed a linear regression model with each component as the only regressor to predict age, sex, and FSIQ, respectively, using the HBN data (Table 1). We found that half of the reproducible components were related to age, FSIQ, and/or sex. Specifically, joint component 2 and components specific to cortical thickness, subcortical GM volume, and mean curvature explained 29%, 18%, 8%, and 3% of total variation in age, respectively, while joint component 1 by itself explained 6% of total variation in FSIQ and predicted sex with good accuracy (AUC=0.79). These results were validated in the NKI-RS cross-sectional sample.

**Table 1:** Predicting age, sex, and FSIQ by individual reproducible component

| Component |  | HBN |  |  |  |  | NKI-RS |  |  |  |  |
| --- | --- | --- | --- | --- | --- | --- | --- | --- | --- | --- | --- |
|  |  | Age |  | FSIQ |  | Sex | Age |  | FSIQ |  | Sex |
|  |  | MAE | R<sub>adj</sub><sup>2</sup> | MAE | R<sub>adj</sub><sup>2</sup> | AUC | MAE | R<sub>adj</sub><sup>2</sup> | MAE | R<sub>adj</sub><sup>2</sup> | AUC |
| *Joint* |  |  |  |  |  |  |  |  |  |  |  |
|  | Comp 1 | 2.70 | 0.01 | 12.60 | **0.06** | **0.79** | 2.87 | 0.00 | 11.69 | **0.05** | **0.80** |
|  | Comp 2 | 2.27 | **0.29** | 13.16 | 0.01 | 0.50 | 3.18 | **0.28** | 11.52 | 0.01 | 0.62 |
| *Specific to travel depth* |  |  |  |  |  |  |  |  |  |  |  |
|  | Comp 1 | 2.70 | **0.01** | 13.18 | 0.00 | 0.52 | 2.87 | **0.01** | 11.83 | 0.01 | 0.53 |
|  | Comp 2 | 2.71 | 0.00 | 13.06 | 0.01 | 0.53 | 2.85 | 0.00 | 11.77 | 0.04 | 0.50 |
| *Specific to* |  |  |  |  |  |  |  |  |  |  |  |
|  | White matter volume | 2.68 | 0.02 | 13.15 | 0.00 | 0.53 | 2.91 | 0.00 | 11.78 | 0.03 | 0.59 |
|  | Cortical thickness | 2.43 | **0.18** | 13.17 | 0.00 | 0.55 | 2.70 | **0.24** | 11.81 | 0.00 | 0.55 |
|  | Gray matter volume | 2.70 | **0.01** | 13.17 | 0.00 | 0.51 | 2.79 | **0.04** | 11.89 | 0.01 | 0.54 |
|  | Mean curvature | 2.67 | **0.03** | 13.14 | 0.00 | 0.51 | 2.88 | **0.03** | 11.86 | 0.01 | 0.54 |
|  | Surface area | 2.70 | **0.01** | 13.15 | 0.00 | 0.50 | 2.83 | **0.02** | 11.88 | 0.00 | 0.51 |
|  | Subcortical gray matter volume | 2.59 | **0.08** | 13.15 | 0.00 | 0.52 | 2.51 | **0.08** | 12.37 | 0.01 | 0.50 |

### 3.2 Brain regions with large loadings in highly reproducible components

To identify what shape measures in which regions could show similar variation patterns across all samples, we examined brain regions and their corresponding shape measures with loading magnitudes greater than 0.1 in the identified components. There are 388 loading estimates in each joint component representing covariation on all seven shape measures across various brain regions. Individual components related to shape measure for cortical and subcortical regions had 62 and 16 loading estimates, respectively. Here, we focused on the two joint components and the component specific to cortical thickness due to their relevance to age, FSIQ, and sex prediction.

A small number of loadings had large magnitudes (defined as loading magnitude greater than 0.1) in both joint component 1 (24 out of 388 loadings, 6.2%) and joint component 2 (27 out of 388 loadings, 7.0%). Specifically, both joint components had large loadings on left and right thalamus proper, ventral diencephalon, and superior frontal WM volumes. Shape measures in those regions predicted age (R_adj_^2^ ranging from 4% to 16%), sex (AUC around 0.70), and FSIQ (R_adj_^2^ ranging from 1% to 3%). Since joint component 1 was largely related to sex and FSIQ while joint component 2 was mostly related to age, brain shape measures with large loadings in both joint components are expected to be related to age, FSIQ, and sex. Regions with large loadings on joint component 1 only (Table 2) were mainly cortical GM volume, surface area, and WM volume in left and right rostral middle frontal and superior frontal, along with GM volume in subcortical regions (caudate, putamen, and hippocampus). Shape measures in those regions individually explained up to 5% of total variation in FSIQ and predicted sex with AUC ranging from 0.6 to 0.74. In contrast, regions with large loadings on joint component 2 only (Table 3) were mostly cortical thickness in the following regions: middle temporal, supramarginal, superior frontal, pars orbitalis, inferior and superior parietal in both hemispheres, left transverse temporal, right lateral orbitofrontal, right rostral anterior cingulate, right precuneus, and right superior temporal. Each shape measure in those regions predicted age (R_adj_^2^ ranging from 3% to 20%). Table 4 summarizes regions with large loadings in the component specific to cortical thickness that by itself explained 18% variation in age. It should be noted that cortical thickness in left and right rostral and caudal anterior cingulate individually explained a decent amount of variation in age (R_adj_^2^ ranging from 5% to 15%).

**Table 2:**
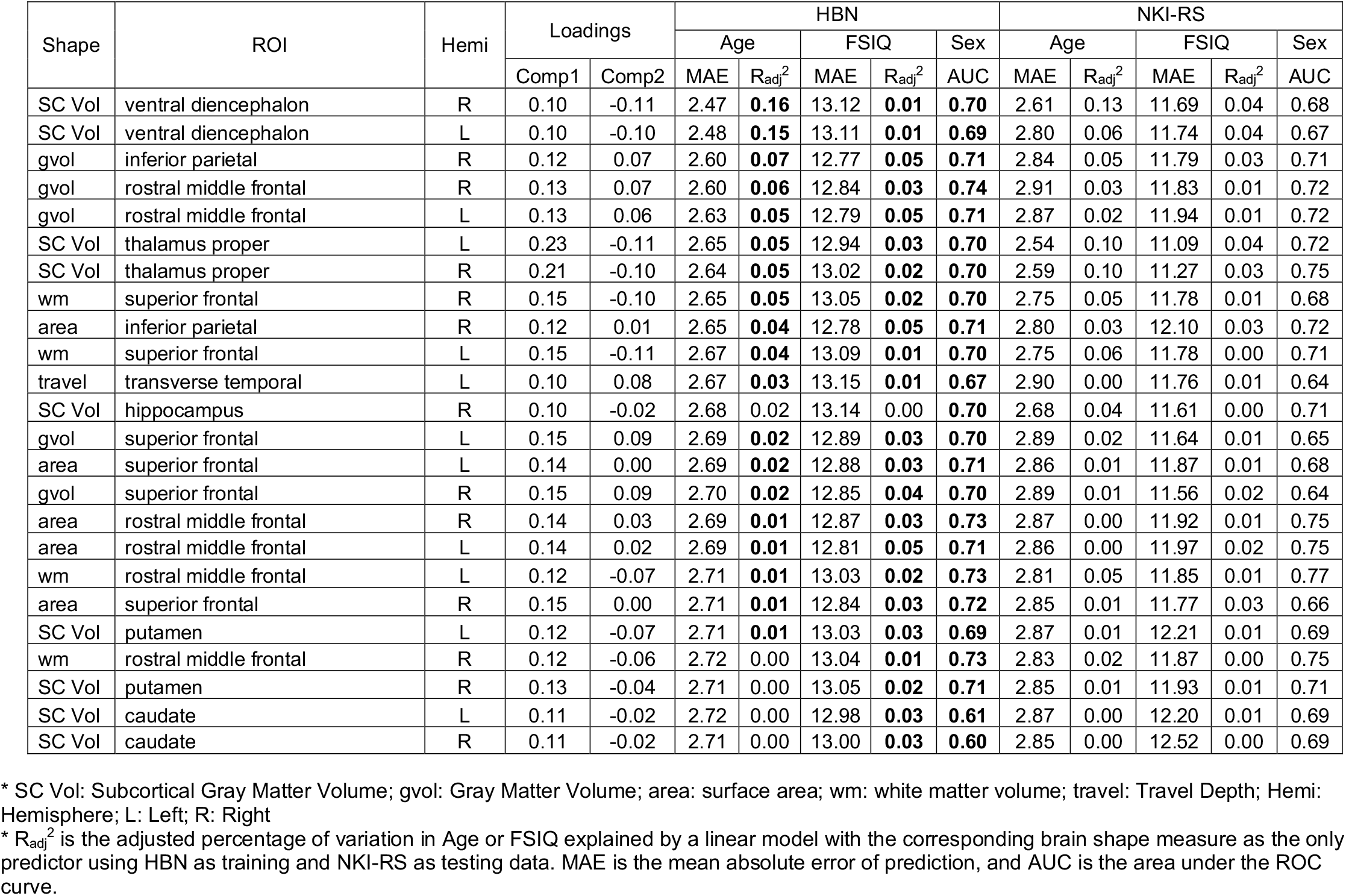
Brain regions with loading magnitudes greater than 0.1 in joint component 1.

**Table 3 Version 2:**
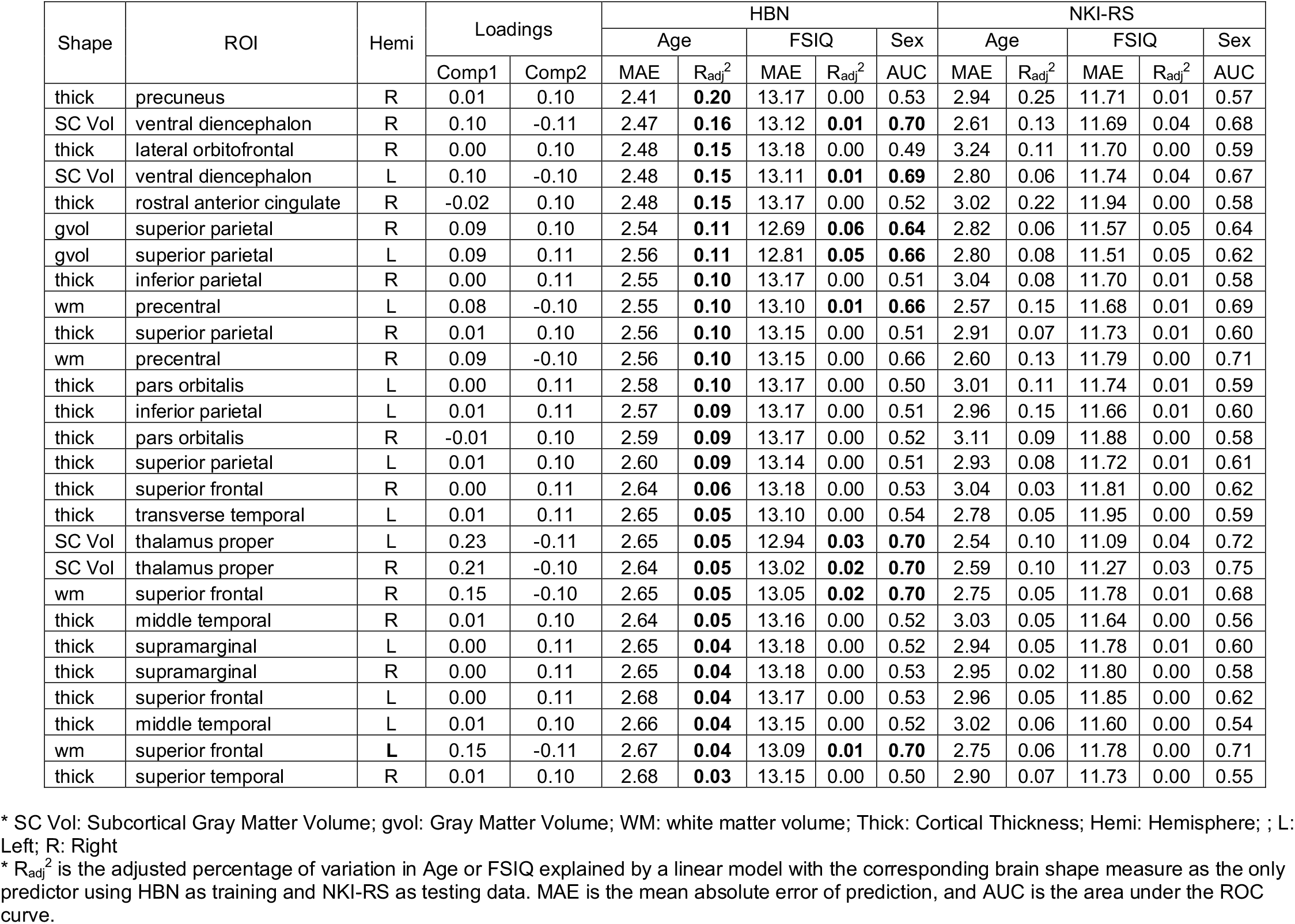
Brain regions with loading magnitudes greater than 0.1 in joint component 2.

| Shape | ROI | Hemi | Loadings |  | HBN |  |  |  |  | NKI-RS |  |  |  |  |
| --- | --- | --- | --- | --- | --- | --- | --- | --- | --- | --- | --- | --- | --- | --- |
|  |  |  |  |  | Age |  | FSIQ |  | Sex | Age |  | FSIQ |  | Sex |
|  |  |  | Comp1 | Comp2 | MAE | R<sub>adj</sub><sup>2</sup> | MAE | R<sub>adj</sub><sup>2</sup> | AUC | MAE | R<sub>adj</sub><sup>2</sup> | MAE | R<sub>adj</sub><sup>2</sup> | AUC |
| thick | precuneus | R | 0.01 | 0.10 | 2.41 | **0.20** | 13.17 | 0.00 | 0.53 | 2.94 | 0.25 | 11.71 | 0.01 | 0.57 |
| SC Vol | ventral diencephalon | R | 0.10 | -0.11 | 2.47 | **0.16** | 13.12 | **0.01** | **0.70** | 2.61 | 0.13 | 11.69 | 0.04 | 0.68 |
| thick | lateral orbitofrontal | R | 0.00 | 0.10 | 2.48 | **0.15** | 13.18 | 0.00 | 0.49 | 3.24 | 0.11 | 11.70 | 0.00 | 0.59 |
| SC Vol | ventral diencephalon | L | 0.10 | -0.10 | 2.48 | **0.15** | 13.11 | **0.01** | **0.69** | 2.80 | 0.06 | 11.74 | 0.04 | 0.67 |
| thick | rostral anterior cingulate | R | -0.02 | 0.10 | 2.48 | **0.15** | 13.17 | 0.00 | 0.52 | 3.02 | 0.22 | 11.94 | 0.00 | 0.58 |
| gvol | superior parietal | R | 0.09 | 0.10 | 2.54 | **0.11** | 12.69 | **0.06** | **0.64** | 2.82 | 0.06 | 11.57 | 0.05 | 0.64 |
| gvol | superior parietal | L | 0.09 | 0.11 | 2.56 | **0.11** | 12.81 | **0.05** | **0.66** | 2.80 | 0.08 | 11.51 | 0.05 | 0.62 |
| thick | inferior parietal | R | 0.00 | 0.11 | 2.55 | **0.10** | 13.17 | 0.00 | 0.51 | 3.04 | 0.08 | 11.70 | 0.01 | 0.58 |
| wm | precentral | L | 0.08 | -0.10 | 2.55 | **0.10** | 13.10 | **0.01** | **0.66** | 2.57 | 0.15 | 11.68 | 0.01 | 0.69 |
| thick | superior parietal | R | 0.01 | 0.10 | 2.56 | **0.10** | 13.15 | 0.00 | 0.51 | 2.91 | 0.07 | 11.73 | 0.01 | 0.60 |
| wm | precentral | R | 0.09 | -0.10 | 2.56 | **0.10** | 13.15 | 0.00 | 0.66 | 2.60 | 0.13 | 11.79 | 0.00 | 0.71 |
| thick | pars orbitalis | L | 0.00 | 0.11 | 2.58 | **0.10** | 13.17 | 0.00 | 0.50 | 3.01 | 0.11 | 11.74 | 0.01 | 0.59 |
| thick | inferior parietal | L | 0.01 | 0.11 | 2.57 | **0.09** | 13.17 | 0.00 | 0.51 | 2.96 | 0.15 | 11.66 | 0.01 | 0.60 |
| thick | pars orbitalis | R | -0.01 | 0.10 | 2.59 | **0.09** | 13.17 | 0.00 | 0.52 | 3.11 | 0.09 | 11.88 | 0.00 | 0.58 |
| thick | superior parietal | L | 0.01 | 0.10 | 2.60 | **0.09** | 13.14 | 0.00 | 0.51 | 2.93 | 0.08 | 11.72 | 0.01 | 0.61 |
| thick | superior frontal | R | 0.00 | 0.11 | 2.64 | **0.06** | 13.18 | 0.00 | 0.53 | 3.04 | 0.03 | 11.81 | 0.00 | 0.62 |
| thick | transverse temporal | L | 0.01 | 0.11 | 2.65 | **0.05** | 13.10 | 0.00 | 0.54 | 2.78 | 0.05 | 11.95 | 0.00 | 0.59 |
| SC Vol | thalamus proper | L | 0.23 | -0.11 | 2.65 | **0.05** | 12.94 | **0.03** | **0.70** | 2.54 | 0.10 | 11.09 | 0.04 | 0.72 |
| SC Vol | thalamus proper | R | 0.21 | -0.10 | 2.64 | **0.05** | 13.02 | **0.02** | **0.70** | 2.59 | 0.10 | 11.27 | 0.03 | 0.75 |
| wm | superior frontal | R | 0.15 | -0.10 | 2.65 | **0.05** | 13.05 | **0.02** | **0.70** | 2.75 | 0.05 | 11.78 | 0.01 | 0.68 |
| thick | middle temporal | R | 0.01 | 0.10 | 2.64 | **0.05** | 13.16 | 0.00 | 0.52 | 3.03 | 0.05 | 11.64 | 0.00 | 0.56 |
| thick | supramarginal | L | 0.00 | 0.11 | 2.65 | **0.04** | 13.18 | 0.00 | 0.52 | 2.94 | 0.05 | 11.78 | 0.01 | 0.60 |
| thick | supramarginal | R | 0.00 | 0.11 | 2.65 | **0.04** | 13.18 | 0.00 | 0.53 | 2.95 | 0.02 | 11.80 | 0.00 | 0.58 |
| thick | superior frontal | L | 0.00 | 0.11 | 2.68 | **0.04** | 13.17 | 0.00 | 0.53 | 2.96 | 0.05 | 11.85 | 0.00 | 0.62 |
| thick | middle temporal | L | 0.01 | 0.10 | 2.66 | **0.04** | 13.15 | 0.00 | 0.52 | 3.02 | 0.06 | 11.60 | 0.00 | 0.54 |
| wm | superior frontal | L | 0.15 | -0.11 | 2.67 | **0.04** | 13.09 | **0.01** | **0.70** | 2.75 | 0.06 | 11.78 | 0.00 | 0.71 |
| thick | superior temporal | R | 0.01 | 0.10 | 2.68 | **0.03** | 13.15 | 0.00 | 0.50 | 2.90 | 0.07 | 11.73 | 0.00 | 0.55 |

\* SC Vol: Subcortical Gray Matter Volume; gvol: Gray Matter Volume; WM: white matter volume; Thick: Cortical Thickness; Hemi: Hemisphere; ; L: Left; R: Right
\* R<sub>adj</sub><sup>2</sup> is the adjusted percentage of variation in Age or FSIQ explained by a linear model with the corresponding brain shape measure as the only predictor using HBN as training and NKI-RS as testing data. MAE is the mean absolute error of prediction, and AUC is the area under the ROC curve.

**Table 4:** Brain regions with loading magnitudes greater than 0.1 in component specific to cortical thickness.

| ROI | Hemi | Loadings | HBN |  |  |  |  | NKI-RS |  |  |  |  |
| --- | --- | --- | --- | --- | --- | --- | --- | --- | --- | --- | --- | --- |
|  |  |  | Age |  | FSIQ |  | Sex | Age |  | FSIQ |  | Sex |
|  |  |  | MAE | R<sub>adj</sub><sup>2</sup> | MAE | R<sub>adj</sub><sup>2</sup> | AUC | MAE | R<sub>adj</sub><sup>2</sup> | MAE | R<sub>adj</sub><sup>2</sup> | AUC |
| rostral anterior cingulate | R | 0.16 | 2.48 | **0.15** | 13.17 | 0.00 | 0.52 | 3.02 | 0.22 | 11.94 | 0.00 | 0.58 |
| rostral anterior cingulate | L | 0.14 | 2.52 | **0.13** | 13.17 | 0.00 | 0.52 | 2.88 | 0.30 | 11.90 | 0.00 | 0.65 |
| caudal anterior cingulate | R | 0.17 | 2.57 | **0.11** | 13.17 | 0.00 | 0.52 | 2.94 | 0.14 | 11.70 | 0.00 | 0.58 |
| inferior parietal | L | -0.10 | 2.57 | **0.09** | 13.17 | 0.00 | 0.51 | 2.96 | 0.15 | 11.66 | 0.01 | 0.60 |
| pars orbitalis | R | -0.11 | 2.59 | **0.09** | 13.17 | 0.00 | 0.52 | 3.11 | 0.09 | 11.88 | 0.00 | 0.58 |
| transverse temporal | L | -0.14 | 2.65 | **0.05** | 13.10 | 0.00 | 0.54 | 2.78 | 0.05 | 11.95 | 0.00 | 0.59 |
| caudal anterior cingulate | L | 0.15 | 2.66 | **0.05** | 13.18 | 0.00 | 0.51 | 2.84 | 0.19 | 11.78 | 0.00 | 0.55 |
| transverse temporal | R | -0.15 | 2.64 | **0.05** | 13.12 | 0.01 | 0.54 | 2.70 | 0.15 | 12.01 | 0.01 | 0.57 |
| supramarginal | L | -0.14 | 2.65 | **0.04** | 13.18 | 0.00 | 0.52 | 2.94 | 0.05 | 11.78 | 0.01 | 0.60 |
| supramarginal | R | -0.14 | 2.65 | **0.04** | 13.18 | 0.00 | 0.53 | 2.95 | 0.02 | 11.80 | 0.00 | 0.58 |
| superior frontal | L | -0.14 | 2.68 | **0.04** | 13.17 | 0.00 | 0.53 | 2.96 | 0.05 | 11.85 | 0.00 | 0.62 |
| middle temporal | L | -0.12 | 2.66 | **0.04** | 13.15 | 0.00 | 0.52 | 3.02 | 0.06 | 11.60 | 0.00 | 0.54 |
| superior temporal | R | -0.14 | 2.68 | **0.03** | 13.15 | 0.00 | 0.50 | 2.90 | 0.07 | 11.73 | 0.00 | 0.55 |
| precentral | L | -0.18 | 2.66 | **0.03** | 13.17 | 0.00 | 0.55 | 2.77 | 0.03 | 11.91 | 0.01 | 0.58 |
| postcentral | R | -0.13 | 2.67 | **0.03** | 13.12 | 0.01 | 0.55 | 2.90 | 0.00 | 11.86 | 0.00 | 0.58 |
| superior temporal | L | -0.18 | 2.70 | **0.02** | 13.17 | 0.00 | 0.51 | 2.92 | 0.01 | 11.81 | 0.00 | 0.56 |
| precentral | R | -0.16 | 2.68 | **0.02** | 13.18 | 0.00 | 0.55 | 2.80 | 0.02 | 11.78 | 0.02 | 0.56 |
| postcentral | L | -0.15 | 2.70 | **0.01** | 13.13 | 0.00 | 0.56 | 2.84 | 0.01 | 11.91 | 0.00 | 0.65 |
| entorhinal | L | -0.44 | 2.70 | **0.01** | 13.17 | 0.00 | 0.52 | 2.83 | 0.04 | 11.85 | 0.00 | 0.52 |
| entorhinal | R | -0.49 | 2.71 | **0.01** | 13.17 | 0.00 | 0.50 | 2.83 | 0.05 | 11.87 | 0.00 | 0.55 |
| caudal middle frontal | R | -0.11 | 2.72 | 0.00 | 13.16 | 0.00 | 0.55 | 2.88 | 0.01 | 11.94 | 0.02 | 0.58 |
| caudal middle frontal | L | -0.14 | 2.72 | 0.00 | 13.17 | 0.00 | 0.54 | 2.86 | 0.00 | 11.85 | 0.00 | 0.56 |

\*Hemi: Hemisphere ; L: Left; R: Right

Our results (Tables 5 – 7) revealed that among all 388 shape measures, those that individually explained at least 10% of variation in age were mainly cortical thickness; those that individually explained at least 4% of variation in FSIQ were either gray matter volume or cortical surface area; and those predicted sex with AUC above 0.70 were a mix of white matter volume, gray matter volume, and surface area.

**Table 5.**
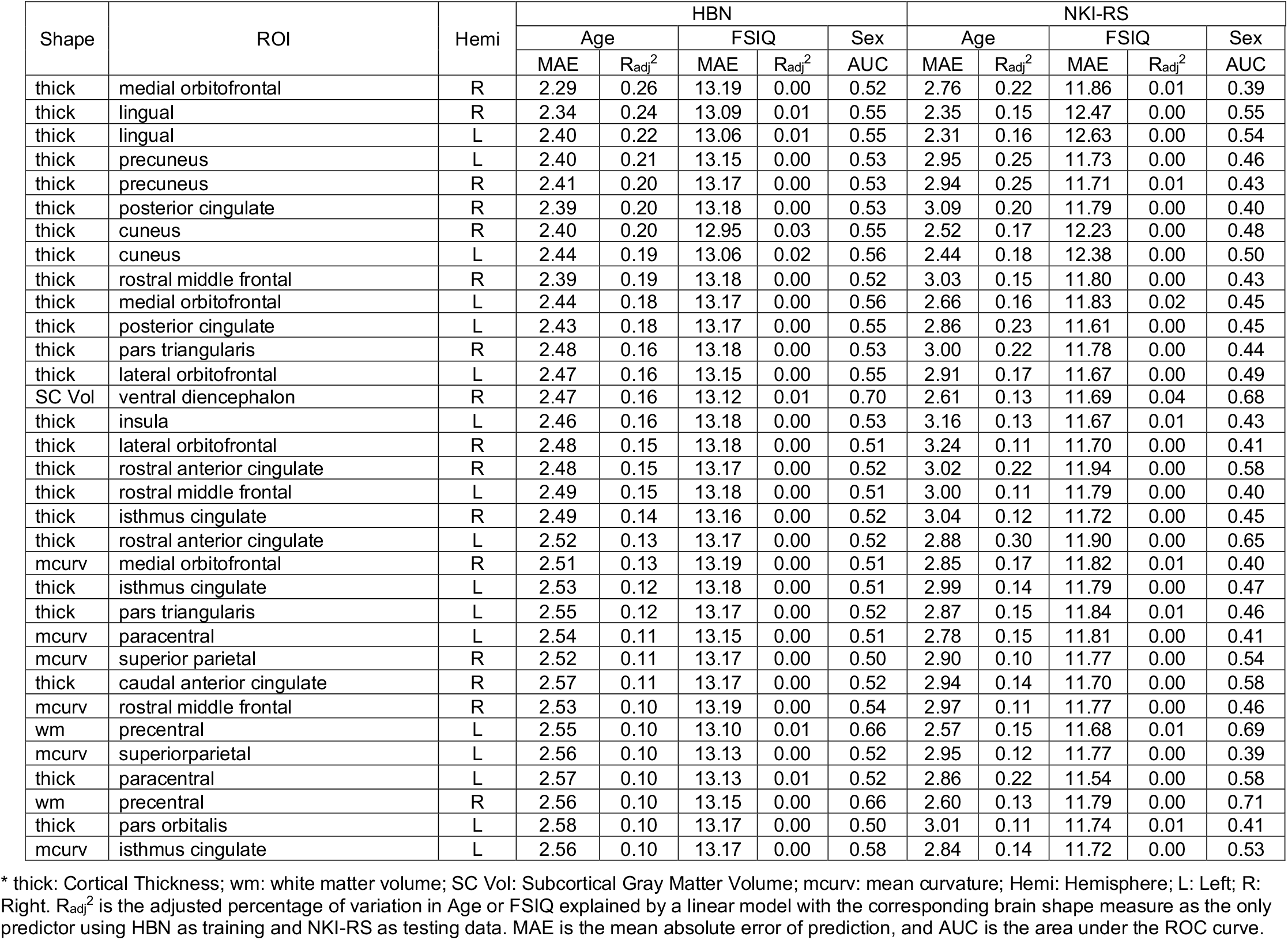
Brain regions predicting age

**Table 6.** Brain regions predicting FSIQ

| Shape | ROI | Hemi | HBN |  |  |  |  | NKI-RS |  |  |  |  |
| --- | --- | --- | --- | --- | --- | --- | --- | --- | --- | --- | --- | --- |
|  |  |  | Age |  | FSIQ |  | Sex | Age |  | FSIQ |  | Sex |
| | | | MAE | $R_{adj}^2$ | MAE | $R_{adj}^2$ | AUC | MAE | $R_{adj}^2$ | MAE | $R_{adj}^2$ | AUC |
| gvol | superior parietal | R | 2.54 | 0.11 | 12.69 | 0.06 | 0.64 | 2.82 | 0.06 | 11.57 | 0.05 | 0.64 |
| area | superior parietal | R | 2.59 | 0.07 | 12.69 | 0.06 | 0.66 | 2.81 | 0.03 | 11.82 | 0.05 | 0.70 |
| gvol | precuneus | R | 2.53 | 0.11 | 12.75 | 0.06 | 0.68 | 2.79 | 0.09 | 11.65 | 0.05 | 0.62 |
| gvol | precuneus | L | 2.54 | 0.10 | 12.71 | 0.06 | 0.70 | 2.80 | 0.08 | 11.60 | 0.06 | 0.66 |
| gvol | middle temporal | R | 2.69 | 0.02 | 12.69 | 0.05 | 0.71 | 2.90 | 0.00 | 11.52 | 0.05 | 0.71 |
| area | precuneus | L | 2.63 | 0.04 | 12.75 | 0.05 | 0.69 | 2.81 | 0.02 | 11.90 | 0.05 | 0.67 |
| area | precuneus | R | 2.60 | 0.06 | 12.82 | 0.05 | 0.69 | 2.78 | 0.04 | 12.00 | 0.04 | 0.63 |
| gvol | superior parietal | L | 2.56 | 0.11 | 12.81 | 0.05 | 0.66 | 2.80 | 0.08 | 11.51 | 0.05 | 0.62 |
| gvol | inferior parietal | R | 2.60 | 0.07 | 12.77 | 0.05 | 0.71 | 2.84 | 0.05 | 11.79 | 0.03 | 0.71 |
| area | superior parietal | L | 2.60 | 0.06 | 12.81 | 0.05 | 0.67 | 2.79 | 0.03 | 11.66 | 0.06 | 0.65 |
| gvol | postcentral | R | 2.60 | 0.07 | 12.77 | 0.05 | 0.64 | 2.90 | 0.03 | 11.37 | 0.04 | 0.61 |
| area | inferior parietal | R | 2.65 | 0.04 | 12.78 | 0.05 | 0.71 | 2.80 | 0.03 | 12.10 | 0.03 | 0.72 |
| gvol | inferior temporal | R | 2.71 | 0.00 | 12.78 | 0.04 | 0.69 | 2.87 | 0.00 | 11.70 | 0.03 | 0.75 |
| area | superior temporal | L | 2.69 | 0.02 | 12.85 | 0.04 | 0.71 | 2.87 | 0.00 | 11.98 | 0.03 | 0.70 |
| area | middle temporal | R | 2.71 | 0.00 | 12.82 | 0.04 | 0.71 | 2.87 | 0.00 | 11.73 | 0.05 | 0.71 |
| area | postcentral | R | 2.61 | 0.06 | 12.82 | 0.04 | 0.69 | 2.88 | 0.03 | 11.42 | 0.04 | 0.64 |
| area | precentral | R | 2.70 | 0.01 | 12.80 | 0.04 | 0.72 | 2.86 | 0.01 | 11.86 | 0.03 | 0.72 |
| gvol | cuneus | R | 2.59 | 0.07 | 12.84 | 0.04 | 0.68 | 2.80 | 0.01 | 12.05 | 0.04 | 0.71 |
| gvol | medial orbitofrontal | R | 2.58 | 0.08 | 12.78 | 0.04 | 0.68 | 2.90 | 0.02 | 11.85 | 0.03 | 0.67 |
| gvol | lateral occipital | L | 2.65 | 0.04 | 12.83 | 0.04 | 0.72 | 2.89 | 0.00 | 11.50 | 0.04 | 0.71 |
| area | lateral occipital | R | 2.69 | 0.02 | 12.88 | 0.04 | 0.74 | 2.91 | 0.01 | 11.49 | 0.07 | 0.75 |
| gvol | middle temporal | L | 2.71 | 0.01 | 12.82 | 0.04 | 0.69 | 2.86 | 0.02 | 11.79 | 0.03 | 0.73 |
| gvol | precentral | R | 2.71 | 0.00 | 12.85 | 0.04 | 0.68 | 2.87 | 0.00 | 11.51 | 0.05 | 0.68 |
| area | lateral occipital | L | 2.69 | 0.02 | 12.84 | 0.04 | 0.72 | 2.87 | 0.00 | 11.48 | 0.05 | 0.74 |
| gvol | fusiform | L | 2.69 | 0.01 | 12.82 | 0.04 | 0.70 | 2.92 | 0.00 | 11.45 | 0.05 | 0.73 |
| area | inferior temporal | R | 2.71 | 0.00 | 12.88 | 0.04 | 0.68 | 2.83 | 0.02 | 11.75 | 0.03 | 0.74 |
| gvol | lateral occipital | R | 2.67 | 0.03 | 12.89 | 0.04 | 0.73 | 2.93 | 0.00 | 11.41 | 0.06 | 0.73 |

\* gvol: Gray Matter Volume; area: surface area; Hemi: Hemisphere; L: Left; R: Right. $R_{adj}^2$ is the adjusted percentage of variation in Age or FSIQ explained by a linear model with the corresponding brain shape measure as the only predictor using HBN as training and NKI-RS as testing data. MAE is the mean absolute error of prediction, and AUC is the area under the ROC curve.

**Table 7.** Brain regions predicting sex

| Shape | ROI | Hemi | HBN |  |  |  |  | NKI-RS |  |  |  |  |
| --- | --- | --- | --- | --- | --- | --- | --- | --- | --- | --- | --- | --- |
|  |  |  | Age |  | FSIQ |  | Sex | Age |  | FSIQ |  | Sex |
|  |  |  | MAE | R<sub>adj</sub><sup>2</sup> | MAE | R<sub>adj</sub><sup>2</sup> | AUC | MAE | R<sub>adj</sub><sup>2</sup> | MAE | R<sub>adj</sub><sup>2</sup> | AUC |
| gvol | insula | R | 2.71 | 0.00 | 12.86 | 0.03 | 0.74 | 2.85 | 0.00 | 11.58 | 0.01 | 0.75 |
| area | lateral occipital | R | 2.69 | 0.02 | 12.88 | 0.04 | 0.74 | 2.91 | 0.01 | 11.49 | 0.07 | 0.75 |
| gvol | fusiform | R | 2.71 | 0.00 | 12.93 | 0.03 | 0.74 | 2.88 | 0.00 | 11.66 | 0.03 | 0.73 |
| gvol | rostral middle frontal | R | 2.60 | 0.06 | 12.84 | 0.03 | 0.74 | 2.91 | 0.03 | 11.83 | 0.01 | 0.72 |
| wm | rostral middle frontal | R | 2.72 | 0.00 | 13.04 | 0.01 | 0.73 | 2.83 | 0.02 | 11.87 | 0.00 | 0.75 |
| wm | inferior temporal | R | 2.71 | 0.01 | 12.88 | 0.03 | 0.73 | 2.79 | 0.06 | 11.52 | 0.04 | 0.76 |
| area | rostral middle frontal | R | 2.69 | 0.01 | 12.87 | 0.03 | 0.73 | 2.87 | 0.00 | 11.92 | 0.01 | 0.75 |
| gvol | lateral occipital | R | 2.67 | 0.03 | 12.89 | 0.04 | 0.73 | 2.93 | 0.00 | 11.41 | 0.06 | 0.73 |
| area | fusiform | R | 2.72 | 0.00 | 13.01 | 0.02 | 0.73 | 2.85 | 0.02 | 11.71 | 0.03 | 0.73 |
| wm | rostral middle frontal | L | 2.71 | 0.01 | 13.03 | 0.02 | 0.73 | 2.81 | 0.05 | 11.85 | 0.01 | 0.77 |
| gvol | insula | L | 2.71 | 0.00 | 12.86 | 0.04 | 0.73 | 2.91 | 0.00 | 11.52 | 0.01 | 0.70 |
| area | precentral | L | 2.69 | 0.01 | 12.79 | 0.04 | 0.73 | 2.88 | 0.01 | 11.71 | 0.02 | 0.70 |
| wm | superior temporal | L | 2.71 | 0.01 | 13.05 | 0.01 | 0.73 | 2.80 | 0.05 | 11.79 | 0.01 | 0.73 |
| area | lateral occipital | L | 2.69 | 0.02 | 12.84 | 0.04 | 0.72 | 2.87 | 0.00 | 11.48 | 0.05 | 0.74 |
| wm | supra marginal | L | 2.71 | 0.00 | 12.98 | 0.02 | 0.72 | 2.82 | 0.05 | 11.79 | 0.01 | 0.77 |
| area | precentral | R | 2.70 | 0.01 | 12.80 | 0.04 | 0.72 | 2.86 | 0.01 | 11.86 | 0.03 | 0.72 |
| gvol | lateral occipital | L | 2.65 | 0.04 | 12.83 | 0.04 | 0.72 | 2.89 | 0.00 | 11.50 | 0.04 | 0.71 |
| gvol | rostral middle frontal | L | 2.63 | 0.05 | 12.79 | 0.05 | 0.71 | 2.87 | 0.02 | 11.94 | 0.01 | 0.72 |
| wm | lateral occipital | R | 2.68 | 0.03 | 13.14 | 0.00 | 0.71 | 2.64 | 0.15 | 11.67 | 0.02 | 0.74 |
| wm | precuneus | L | 2.71 | 0.01 | 12.93 | 0.03 | 0.71 | 2.78 | 0.06 | 11.65 | 0.03 | 0.75 |
| area | rostral middle frontal | L | 2.69 | 0.01 | 12.81 | 0.05 | 0.71 | 2.86 | 0.00 | 11.97 | 0.02 | 0.75 |
| wm | inferior parietal | R | 2.71 | 0.01 | 13.00 | 0.02 | 0.71 | 2.83 | 0.03 | 11.86 | 0.03 | 0.77 |
| wm | pars triangularis | L | 2.71 | 0.00 | 13.06 | 0.01 | 0.71 | 2.82 | 0.02 | 11.60 | 0.01 | 0.72 |
| gvol | isthmus cingulate | L | 2.64 | 0.04 | 12.99 | 0.02 | 0.71 | 2.86 | 0.00 | 11.91 | 0.01 | 0.74 |
| wm | lateral orbitofrontal | R | 2.68 | 0.02 | 13.05 | 0.01 | 0.71 | 2.71 | 0.08 | 11.48 | 0.04 | 0.70 |
| area | supramarginal | L | 2.65 | 0.04 | 12.83 | 0.03 | 0.71 | 2.84 | 0.00 | 11.98 | 0.02 | 0.70 |
| area | middle temporal | R | 2.71 | 0.00 | 12.82 | 0.04 | 0.71 | 2.87 | 0.00 | 11.73 | 0.05 | 0.71 |
| gvol | inferior parietal | R | 2.60 | 0.07 | 12.77 | 0.05 | 0.71 | 2.84 | 0.05 | 11.79 | 0.03 | 0.71 |
| wm | inferior temporal | L | 2.69 | 0.02 | 12.96 | 0.02 | 0.71 | 2.77 | 0.04 | 11.44 | 0.05 | 0.78 |
| area | superior temporal | L | 2.69 | 0.02 | 12.85 | 0.04 | 0.71 | 2.87 | 0.00 | 11.98 | 0.03 | 0.70 |
| area | inferior parietal | R | 2.65 | 0.04 | 12.78 | 0.05 | 0.71 | 2.80 | 0.03 | 12.10 | 0.03 | 0.72 |
| SC Vol | putamen | R | 2.71 | 0.00 | 13.05 | 0.02 | 0.71 | 2.85 | 0.01 | 11.93 | 0.01 | 0.71 |
| wm | lateral orbitofrontal | L | 2.65 | 0.05 | 13.06 | 0.01 | 0.71 | 2.63 | 0.12 | 11.53 | 0.02 | 0.73 |
| gvol | middle temporal | R | 2.69 | 0.02 | 12.69 | 0.05 | 0.71 | 2.90 | 0.00 | 11.52 | 0.05 | 0.71 |

\* wm: white matter volume; gvol: Gray Matter Volume; area: surface area; SC Vol: Subcortical Gray Matter Volume; Hemi: Hemisphere; L: Left; R: Right. R<sub>adj</sub><sup>2</sup> is the adjusted percentage of variation in Age or FSIQ explained by a linear model with the corresponding brain shape measure as the predictor using HBN as training and NKI-RS as testing data. MAE is the mean absolute error of prediction, and AUC is the area under the ROC curve.

### 3.3. Predicting age, sex, and FSIQ

Next, we aimed to determine the optimal multi-feature prediction models for age, sex and FSIQ. Analyses showed that the 35 joint and individual components which we had identified in HBN data could accurately predict age. Table 8 summarizes model prediction accuracy measures from all seven sets of input predictors. The model with total volumes and joint and individual components gave the best prediction accuracy in term of age (MAE=1.41 years, R_adj_^2^=0.71) using the model trained by the HBN data and making prediction in the independent NKI-RS cross-sectional dataset. Prediction accuracy of the same model predicting the HBN data was nearly identical (MAE=1.42 years, R_adj_^2^=0.70). We note that the joint components and the individual components each made unique contributions in predicting age. Thus, while the joint and independent components together explained 67% of total variation in age, models with the joint components only and with the individual components only explained 31% and 35% of total variation in age, respectively. Although the model with total volumes as the only predictors explained 34% of variation in age, total volumes explained only 3% of variation in age after controlling for the joint and individual components in the model.

**Table 8:**
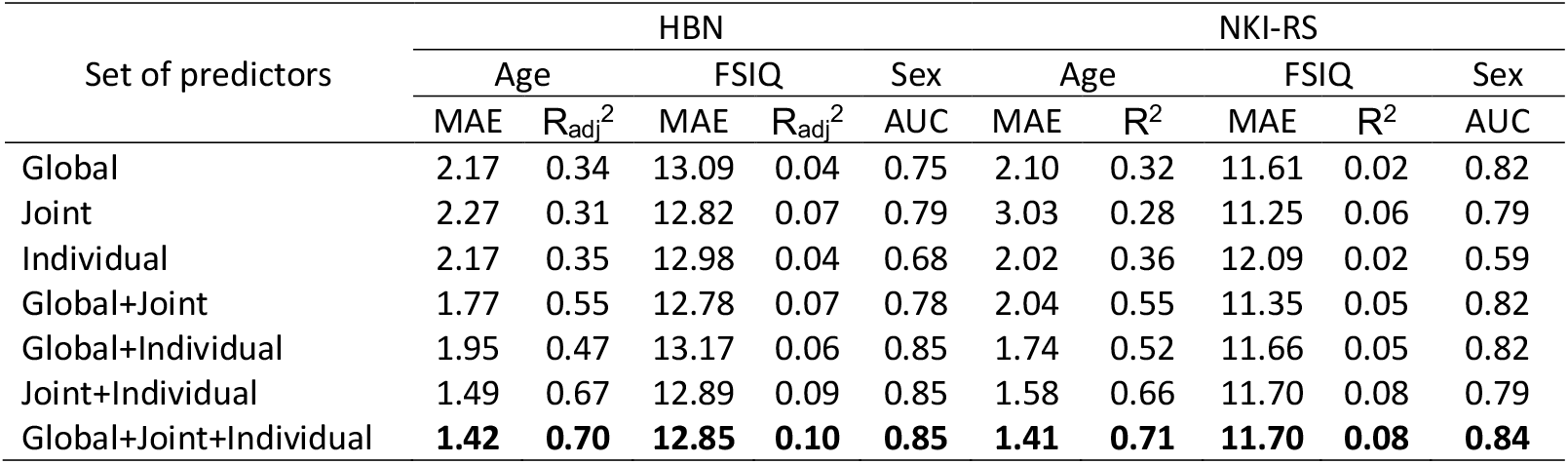
Model prediction accuracy by Ridge Regression using HBN as the training data and NKI-RS as the testing data. Models with best prediction accuracy are in bold.

The model with the best prediction accuracy for sex (AUC=0.85) and FSIQ (MAE=12.85 points, R_adj_^2^=0.10) in the HBN data was the one with joint and individual components plus total volumes, and the results were validated in the independent NKI-RS data for both sex (AUC=0.84) and FSIQ (MAE=11.70, R^2^=0.08). Similar to age prediction, the joint components and the individual components made separate contributions in predicting FSIQ. Interestingly, we observed that the age prediction residuals, defined as chronological age - predicted age, were negatively associated with FSIQ (*r* = −0.11, p-value < 0.001).

### 3.4. Validation by test-retest and longitudinal data

Our model has excellent test-retest reliability. The joint and individual component scores based on the JIVE analysis of the HBN data were calculated for both test and retest data. The predicted brain age and FSIQ were obtained for both test and retest subjects using the prediction models from ridge regression analysis of the HBN data. The intraclass correlation coefficient for measuring agreement of the predicted values between test and retest subjects was 0.97 for age and 0.96 for FSIQ.

Finally, for the longitudinal data available in NKI-RS, we noted that the longitudinal interval between scans was somewhat variable (mean = 1.22 year later ± 0.29). As such, we took the opportunity to test the ability to detect differences in this interval across subjects. We found that the median and median absolute deviation of absolute differences between the real age difference and the predicted age difference were 0.53 and 0.47 years, respectively. A significant positive correlation (*r* = 0.24, p-value =0.04) was observed between the real age difference and the predicted age difference.

## 4. Discussion

### 4.1 Integrative analysis of disparate data types

A major challenge in the analysis of brain imaging studies is the integration of disparate data types, given the substantial covariation across different shape measures as well as within each shape measure. If such covariation in brain measurements cannot be effectively summarized into a lower dimensional representation of brain structure, then no satisfactory performance can be produced by existing statistical models (Bzdok and Yeo, 2017). This highlights the importance of dimension reduction in analysis of neuroimaging data (Cunningham and Yu, 2014; Zhao and Castellanos, 2016).

Motivated by this, we have extended a novel dimension reduction method, JIVE (Lock et al., 2013), initially developed for integrative analysis of different genetic data types, to brain imaging data analysis. We selected JIVE for two main reasons: simultaneous detection of covariation among different data types, and its ability to extract interpretable brain features. To our knowledge, no prior work has attempted to systematically study covariation patterns and changes on such a comprehensive set of brain structure measures across early developmental ages. Extracting interpretable brain features is particularly important in brain imaging studies where imaging confounds or artifacts, such as scanner differences, can easily dominate the results. JIVE is an extension of Principal Component Analysis for extracting linear features (i.e., components) within data. Therefore, it has the advantage of mapping each component back to the original variables and thus making interpretation possible. Our study shows that JIVE can effectively identify meaningful and highly reproducible common and distinct variations across a number of brain cortical and subcortical structural measures that predict age, sex, and FSIQ.

### 4.2 Highly reproducible common and distinct variations in brain shape measures

Prior research on identifying structural covariation has been primarily focused on either a single structural measure (Mechelli et al., 2005), or on analyzing different structural measures separately (Remer et al., 2017; Wierenga et al., 2014). Individual analyses might fail to capture the critical associations between different structural measures (Lock et al., 2013). In this study, we explored simultaneously the covariation patterns across a range of structural measures and all brain regions examined. We demonstrated covariation patterns that exist across different brain cortical and subcortical structure measures in various brain regions, as well as those that are specific to individual brain shape measures. Joint covariation patterns common to all shape measures and a subset of individual covariation patterns specific to travel depth, subcortical gray matter volume, mean curvature, and cortical thickness were remarkably consistent between HBN and NKI-RS datasets.

A question of interest is whether the reproducible covariation patterns in the present investigation are of biological interest. Interpreting the extracted covariation patterns was challenging and not straight-forward. We may not be able to specify the functionality of each identified covariation pattern, which is partly due to the functionality of different brain cortical and subcortical regions not yet being fully characterized (Mechelli et al., 2005.) As a starting point, we focused on their relevance to age, sex, and FSIQ prediction. We found that joint component 2 by itself explained 29% of total variation in age and that almost all its loadings with magnitude greater than 0.1 were cortical thickness measures mainly from frontal and temporal lobes. The component specific to cortical thickness by itself explained 18% of total variation in age. In addition, most of the shape measures in individual brain regions that explained at least 10% of total variation in age were cortical thickness (Table 5). Taken together, these results highlight the importance of cortical thickness as a sensitive index of brain development (Dosenbach et al., 2010; Khundrakpam et al., 2015). In contrast, joint component 1 was not related to age, but predicted sex with AUC of 0.79 by itself. Joint component 1 confirmed that sex differences are most extreme in subcortical volumes (Ritchie et al., 2018). It also revealed sex differences in three shape measures (white matter volume, gray matter volume, and surface area) in superior frontal and rostral middle frontal regions – two primary dorsolateral prefrontal cortex (dlPFC) regions. Lastly, joint component 1 explained about 6% of total variation in FSIQ. We observed that FSIQ correlated mostly with shape measures in the dlPFC regions and thalamus proper. Covariation between dlPFC regions and thalamus proper is consistent with the thalamus as a key subcortical relay underlying executive functions, in line with findings from a tractography study (Le Reste et al., 2016). Therefore, although the neurobiological properties for the cortical and subcortical covariation patterns we identified remain to be fully dissected, these covariations can serve as a starting point for characterizing patterns related to brain maturation and cognitive development.

The covariation patterns specific to GM volume, cortical thickness, and mean curvature were in general less consistent between HBN and NKI-RS datasets (Figure 2). One possible explanation is that developmental trajectories of those measures change across different stages of life and their covariation patterns might be heterogeneous and nonlinear. For example, various studies have shown that cortical thickness exhibits an inverted-U trajectory from childhood to adulthood, GM declines in a regionally heterogeneous pattern, and mean curvature follows a combination of linear, quadratic, and logarithmic trajectories (Brain Development Cooperative, 2012; Giedd et al., 1999; Remer et al., 2017; Shaw et al., 2008; Tamnes et al., 2017; Wierenga et al., 2014). Therefore, JIVE, a method mainly used to identify linear covariation patterns, might fail to recognize nonlinear covariation patterns among shape measures. Furthermore, distinct genetic influences on surface area and thickness (Chen et al., 2013; Panizzon et al., 2009) and different evolutionary processes (Geschwind and Rakic, 2013; Stiles and Jernigan, 2010) may introduce complex individual variability, making it difficult to characterize the covariation patterns of those shape measures.

### 4.3 Age, sex and FSIQ prediction

One of the aims of the current study was to test whether common structural covariations and those which are specific within each shape measure can predict biological factors such as age, sex, and FSIQ. Ridge regression models with the identified brain covariation patterns as predictors accurately predicted age. The model with JIVE joint and individual components along with total volumes produced a mean absolute age prediction error of 1.41 years and explained 71% of total variation in age, making it one of the best age prediction models in the recent literature (Brown et al., 2012; Erus et al., 2015; Franke et al., 2012; Khundrakpam et al., 2015; Lewis et al., 2018). We also found that the JIVE joint and individual components together predicted sex with high accuracy (AUC=0.85) and explained 10% of total variation in FSIQ, achieving better prediction results than a recent study using T1 white/gray contrast to predict FSIQ (Lewis et al., 2018). To test the validity of the model’s predictive performance, we applied it to an independent dataset (NKI-RS) and the results showed that the model generalized to the new dataset, has excellent test-retest reliability, and was able to capture longitudinal changes.

Finally, there is a growing interest in relating individual differences in brain maturation to cognition (Burgaleta et al., 2014; Erus et al., 2015; Lewis et al., 2018; Shaw et al., 2006). For example, (Erus et al., 2015) and (Lewis et al., 2018) both used a residual-based approach to investigate the relationship between brain development and cognition, where brain development was approximated by the age prediction residuals. Both studies showed a significant relationship between brain development and cognition, but the direction of this relationship was opposite in the two studies. Interestingly, we also found a significant negative correlation between the age prediction residuals and FSIQ, indicating that individuals with brain-based age younger than their chronological age had lower FSIQ. This aligns well with the finding of Erus et al. (Erus et al., 2015), although they assessed cognitive test performance in several domains rather than FSIQ. Nevertheless, we note that prediction residuals explained only 1.3% of total variation in FSIQ, so the statistically significant association between age prediction residuals and FSIQ we observed are not yet clinically important. In addition, interpretation of these associations should be made with caution, as conditions such as diabetes, schizophrenia, and traumatic brain injury have been linked to faster brain aging (Cole et al., 2015; Franke et al., 2013). Still, the directionality of our finding has face validity and it warrants further investigation.

### 4.4. Reproducibility

An important aspect of the present work is the high bar employed for examining reproducibility. The JIVE components and prediction models were estimated using a sample that was entirely distinct from that used for testing. This is in contrast to combining the samples and using a less stringent cross-validation strategy or developing unique estimates for each sample. The two samples differed with respect to recruitment strategy, study design, the scanners and imaging protocols used; similarities included the principal investigator. We believe future work would merit from raising the bar for reproducibility to such standards as large-scale samples emerge.

### 4.5 Limitations

Studies have reported that cortical structure development is characterized in general by a mix of linear, curvilinear, and quadratic trajectories with region-specific developmental variations (Remer et al., 2017; Tamnes et al., 2017; Wierenga et al., 2014). However, JIVE is limited to detecting linear covariation patterns. Therefore, we were not able to detect nonlinear covariation patterns that may be common and distinct across cortical shape measures.

A second limitation is the cross-sectional nature of most of the data we examined. Covariation patterns captured by JIVE analysis do not account for within-participant variation. Brain structure changes dynamically throughout the lifespan and the rate of change varies by brain structure measures and brain regions (Storsve et al., 2014). Future explorations should include applying this analytical framework to longitudinal studies, evaluating how changes in various brain structure measures covary and whether the covariation patterns are related to developmental behaviors.

Third, there are concerns about the representativeness of the datasets employed. While the NKI-RS is a community-ascertained sample, with 42.4% having mental health disorders, the HBN is a self-referred community sample, with 81.5% of individuals having one or more mental health or learning diagnoses. Given the relatively large difference in effect sizes observed between age effects and those of psychiatric disorders, it is not surprising that performance was high despite the phenotypic variation in the HBN samples. Nonetheless, the heterogeneity may have hindered our ability to optimally detect small effects, such as IQ. As large-scale, representative samples such as NIH ABCD emerge, we expect that there will be opportunities to develop more optimized, representative models.

Lastly, the brain parcellation map used in this study was based on the Desikan–Killiany–Tourville labeling protocol (Klein and Tourville, 2012) which consists of 62 regions. Khundrakpam et al. (Khundrakpam et al., 2015) reported that the spatial scale of brain parcellation increases the accuracy of age prediction by cortical thickness, with the best estimations obtained for spatial resolutions consisting of 2,560 and 10,240 brain parcels. It is likely that JIVE analysis based on data from brain labels at finer resolutions such as HCP-MMP (Glasser et al., 2016) might improve the model prediction accuracy and produce structural covariation patterns with increased biological interpretability.

In summary, we used JIVE to identify the consistent covariation patterns common across as well as specific to different cortical and subcortical shape measures. These covariation patterns can accurately predict age and sex, and they also provide better prediction for FSIQ than the existing literature, while retaining neurobiological interpretability. Most importantly, we validated our results in an independent dataset, suggesting the generalizability of our approach in brain imaging studies.

#### Declarations of interest

None

## Acknowledgements

This research was in part supported by National Institute of Mental Health Grant U01MH099059 and R01MH091864 to M.P.M. and the endowment provided by Phyllis Green and Randolph Cōwen. A.K. received funding from the following grants: National Institute of Mental Health R01 MH084029, U01 MH074813, and U01 supplement 3U01MH092250-03S1, and the National Institute on Alcohol Abuse and Alcoholism NCANDA-USA Consortium BD2K supplement. The funders had no role in study design, data collection and analysis, decision to publish, or preparation of the manuscript.

